# Transduction of gluconeogenic enzymes prolongs cone photoreceptor survival and function in models of retinitis pigmentosa

**DOI:** 10.1101/569665

**Authors:** Yashodhan Chinchore, Tedi Begaj, Christelle Guillermeir, Matthew L. Steinhauser, Claudio Punzo, Constance L. Cepko

**Affiliations:** Department of Genetics, Department of Ophthalmology, Harvard Medical School, Boston, MA; Center for Nanoimaging, Division of Genetics, Brigham and Women’s Hospital, Boston, MA; Howard Hughes Medical Institute; Department of Medicine, Harvard Medical School, Boston, MA; Department of Ophthalmology and Visual Sciences, University of Massachusetts Medical School, Worcester, MA

## Abstract

The hereditary nature of many retinal degenerative disorders makes them potentially amenable to corrective gene therapies. Numerous clinical trials are ongoing with the goal to rectify the genetic defect in the afflicted cell types. However, the personalized nature of these approaches excludes many patients for whom the underlying mutation is not mapped, or the number of affected individuals is too few to develop a commercially viable therapy (*vide infra*). Thus, a therapy that can delay visual impairment irrespective of the underlying genetic etiology can satisfy this unmet medical need. Here, we demonstrate the utility of such an approach in retinitis pigmentosa (RP) by promoting survival of cone photoreceptors by targeting metabolic stress. These cells are not primarily affected by the inherited mutation, but their non-autonomous demise leads to a decline in daylight vision, greatly reducing the quality of life. We designed adeno-associated virus (AAV) vectors that promote gluconeogenesis- a pathway found in the liver which produces glucose in response to hypoglycemia. Retinal transduction with these vectors resulted in improved cone survival and delayed a decline in visual acuity in three different RP mouse models. Because this approach extended visual function independent of the primary mutation, therapies emanating from this approach could be used as a treatment option for a genetically heterogenous cohort of patients.

## Introduction

Adeno-associated virus (AAV)-mediated gene therapy is an effective means of correcting genetic deficits, particularly for ocular diseases (Auricchio et al., 2017). Its recent approval as a treatment for Leber’s congenital amaurosis 2 (LCA2) (2018), a blinding disease, highlights its safety and efficacy when delivered to the eye. This treatment, and several others, correct a deficit by supplying a wild type copy of a defective gene. This single gene approach will likely be limited to those diseases with a significant number of affected individuals, owing to the expense and difficulties associated with clinical trials. Diseases that lead to loss of vision are due to mutations in a large number of genes, >250 (Duncan et al., 2018;RetNet), and often affect a very small number of individuals. In addition, complementation-based approaches can fall short when gene dosage has to be fine-tuned, and/or a genetic etiology is unknown. A gene-independent strategy to prolong vision could thus be an effective way to broaden the reach of gene therapy while sidestepping the aforementioned hurdles.

Retinitis pigmentosa (RP) is a disease where vision is lost due to the dysfunction, often followed by death, of photoreceptor cells. It is characterized by loss of dim light vision in early stages followed by progressive deterioration of peripheral, daylight and color vision. Mapping studies have uncovered ~80 genetic loci (Ran et al., 2014; RetNet), enabling the diagnostic tests for some patients. However, for many patients the underlying mutations remain unmapped, making the diagnosis and a resulting “corrective” treatment, a challenging task. The majority of RP disease genes primarily or exclusively affect the function of rod photoreceptors-cells that mediate achromatic and low light vision. The affected rods eventually die, leading to dysfunction and loss of cones, which mediate daylight vision. The effects on cones are thus due to non-autonomous causes, as cones typically do not express the disease gene. This loss of cone function coincides with the decline in quality of life of the RP patients and can potentially be targeted to save the daylight vision regardless of the primary mutation that affects the rods. Proposed causes of cone death include loss of retinal architecture (Roger et al., 2012), trophic support (Mohand-Said et al., 1998; Steinberg, 1994), toxin release by the dying cells (Ulshafer et al., 1990), microglial activation (Silverman and Wong, 2018), oxidative stress (Komeima et al., 2006) and metabolic perturbation (Punzo et al., 2009).

Several lines of evidence suggest that cones suffer from metabolic dysfunction in RP. First, cones in four different RP mouse models display a reduction in mTORC1 signaling (Punzo et al., 2009) and activation of this pathway can prolong cone survival (Venkatesh et al., 2015). The stressed cones upregulate chaperone-mediated autophagy and their survival can be enhanced by systemic treatment with insulin (Punzo et al., 2009). In addition, other genetic approaches that target photoreceptor metabolism have also been shown to increase survival (Zhang et al., 2016). The active state of mTOR in WT cones was found to be sensitive to extracellular glucose (Punzo et al., 2009), leading to the suggestion that glucose levels or glucose metabolism might prove limiting after rod death. The shortage of glucose to cones, after loss of glucose-consuming rods is counterintuitive and the mechanisms that might cause such limitations are unclear, but there is some evidence for two models. One scenario is that there is reduced uptake of glucose by cones as a result of reduced availability of rod-derived cone viability factor (RDCVF). RdCVF indirectly stabilizes a glucose transporter on the cone plasma membrane (Aït-Ali et al., 2015) and its deficiency might limit glucose uptake by reducing functional transporter on the cone surface. Another scenario is that there is sequestration of glucose in the retinal pigmented epithelium (RPE) after rod death (Wang et al., 2016) thereby preventing delivery of glucose from the choriocapillaris. As there are no direct measures of intracellular glucose levels in cones in WT and diseased states, it has been difficult to assess these models.

Since it is unclear what the mechanisms might be that limit uptake of glucose by cones (Aït-Ali et al., 2015) and/or delivery of glucose to cones (Wang et al., 2016), we hypothesized that enabling cells to synthesize their own glucose might bypass these limitations. Gluconeogenesis is such a process where glucose can be synthesized from several different carbon sources (Exton, 1972). We thus devised AAV vectors that deliver gluconeogenic enzymes to RP cones. We found that these vectors led to improved cone survival and function. In addition, we tested whether the gluconeogenic AAVs might provide an additive benefit to that of Nrf2. We previously showed that AAV-mediated delivery of Nrf2, a transcription factor that regulates many genes that counter oxidative damage and inflammation, prolonged cone survival and function. These two treatments were additive for cone survival.

## Results

We devised a therapy using delivery of gluconeogenic enzymes, based on the rationale described above, as well as two additional observations made in our studies of metabolism of photoreceptors of wild type and RP mice. We used multi-isotope imaging mass spectrometry (MIMS)(Steinhauser et al., 2012) to track the fate of carbon derived from ^13^C-labelled glucose, and nitrogen derived from ^15^N-labelled glutamine. An outcome of tissue processing for this method is that signals arise solely from isotopic incorporation in the fixable biomass, while soluble, low-molecular weight metabolites are leached out in the wash steps. Radiolabelled amino acids have been used previously to demonstrate their incorporation during outer segment biogenesis (Young, 1967). Those data served as a benchmark to examine the distribution of ^15^N-Glutamine, especially in organelles for which anabolism is essential, and to assess the ^13^C-Glucose-derived carbon colocalization. In RP, the loss of cone outer segments precedes cell death. Intraperitoneal injection of ^13^C-glucose and ^15^N-glutamine into wild type mice was carried out 36 and 12 hours before sacrifice. Retinal sections were imaged by MIMS, which provides high-resolution (<100 nm) images of cellular structures coupled with quantitative mapping of stable isotope tracers. Mass images based on the intensity of CN^−^ and P^−^ delineated stereotypical aspects of retinal histology and guided the extraction of quantitative labeling data. Incorporation of stable isotope tracers (^13^C and ^15^N) was quantified by an increase in the respective isotope ratios above the natural background. First, we observed that, in wild type photoreceptors, glucose-derived carbons were incorporated into the outer segments (OS) (Figure 1A), with signals detectable throughout this structure. The glutamine label from the injected pulse was preferentially localized at the base of the OS (Figure 1A). Interestingly, the puncta corresponding to the carbon signal displayed a significantly high degree of colocalization with glutamine-derived nitrogen puncta (Figure 1B). This observation further supports a biosynthetic route taken by some of the glucose-derived carbons, as opposed to complete oxidation and release as carbon dioxide or secretion as lactate. We confirmed these observations by using ^2^H-labelled glucose and ^15^N-glutamine in wild type mice (Supplementary figure 1A). Again, we observed intense labeling of ^15^N-positive puncta at the base of the OS. The ^2^H signal from glucose displayed a significantly high degree of colocalization with labelled glutamine from the injected label (Supplementary figure 1B).

The intensity of incorporated glucose-derived ^13^C with the ^15^N was compared across cell types and subcellular structures of the eye (Supplementary figure 1C). While we observed a positive linear relationship between biomass-incorporated glucose and glutamine, certain cell types (such as the RPE) and cellular structures (such as the OS and INL) clustered on distinct sides of the regression line. We also assessed the ratio of ^13^C/^15^N to determine if there was a preference to use glucose over glutamine for anabolic reactions (Supplementary figure 1D). We noted significantly elevated ratios in photoreceptor nuclei and the OS compared to the inner segments, and other cell types. These intercellular and intracellular differences might reflect metabolic compartmentalization arising due to local metabolite availability and/or flux (Hurley et al., 2015).

We also examined ^13^C-glucose and ^15^N-glutamine labelling in 5-week old *rd*^*1*^ mice (Supplementary figure 2A). At this stage in *rd*^*1*^ mice, rod death is complete, and thus the only photoreceptors in the ONL are the cones (which do not express the mutant gene). These cones are devoid of the OS. Thus, we quantified the ^13^C and ^15^N incorporation in the nuclei, and these values were used for unpaired comparison with values obtained from adjacent RPE cells and nuclei in the INL (Supplementary figure 2B). Though there was no significant difference between the mean ^13^C/^15^N ratio values between ONL and INL, which would be expected if only cones underwent glucose deprivation, we noted a marked decline in mean value of this ratio in the *rd*^*1*^ ONL compared to the wild type ONL (Supplementary figure 1D). Interestingly, there was no detectable difference in mean raw ^13^C incorporation values between the ONL and the INL in the *rd*^*1*^ wild type ONL (Supplementary figure 1E).

**Figure 1:**
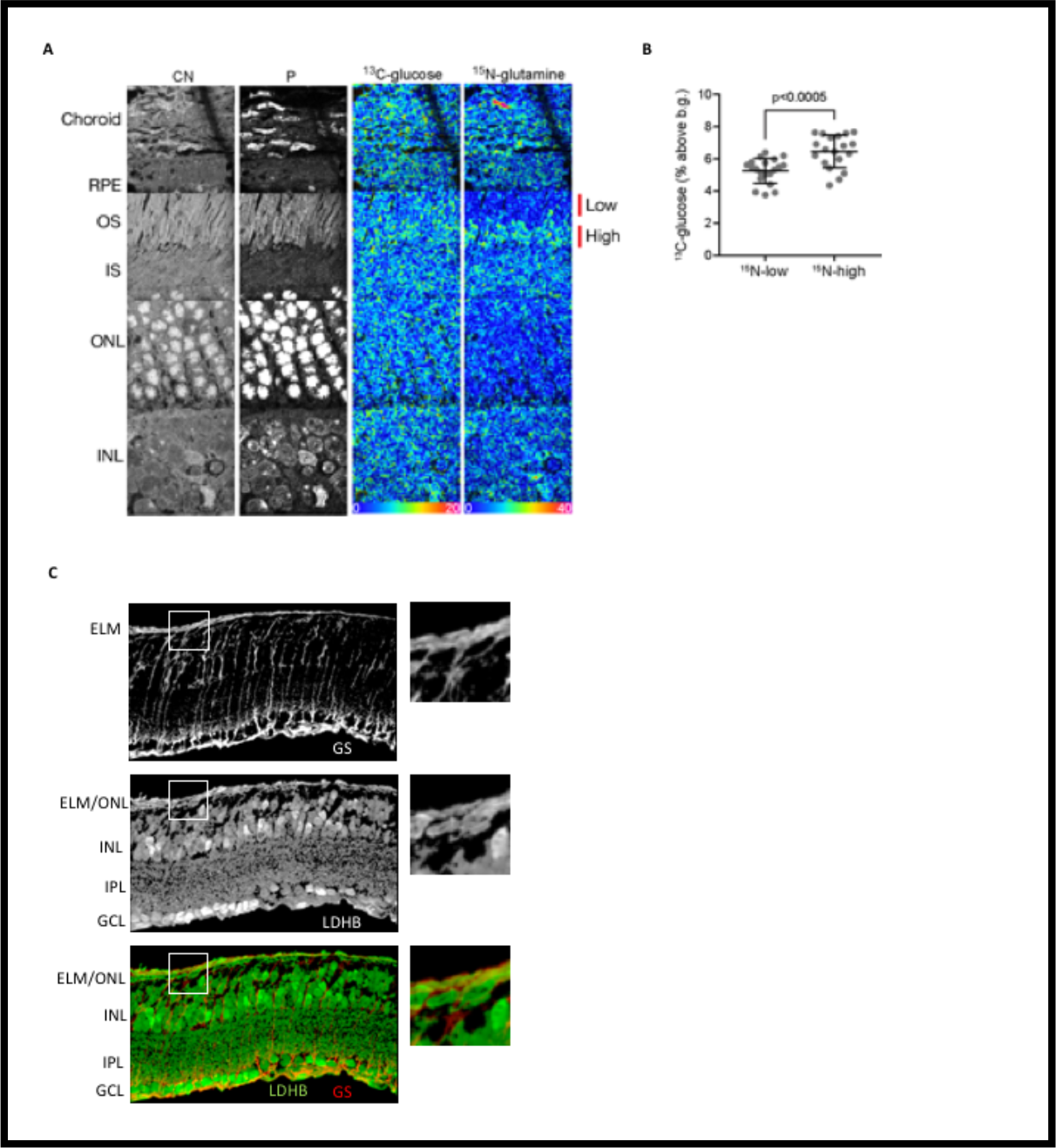
Metabolic rationale for gluconeogenesis. (A). MIMS imaging of an eye from post-natal day 35 CD-1 (wild type) animal injected with ^13^C-glucose and ^15^N-glutamine. Images derived from intensities of ^12^C^14^N (CN) and ^31^P (P) reveal stereotypical histological features of the eye. Incorporation of ^13^C-glucose and ^15^N-glutamine is shown visually by hue saturation intensity (HSI) ratio images (^15^N: ^12^C^15^N/^12^C^14^N and 13C: ^13^C^14^N/^12^C^14^N). In these HIS images, the blue end of the rainbow scale is set to the natural background (0) and the upper bound of the scale is set at a % above natural background that reveals labelling differences. Changes in scaling do not affect the underlying quantitative data as extracted in “B.” Glutamine-derived label was enriched in the proximal aspect of outer segment region (“High”). OS, photoreceptor outer segments, IS, photoreceptor inner segments, ONL, outer nuclear layer, OPL, outer, plexiform layer, INL, inner nuclear layer. (B) Quantification of ^13^C-glucose incorporation in regions of “High” or “Low” ^15^N-glutamine labelling. Individual region-of-interest (ROI) were selected with lateral dimensions not exceeding each OS width. A single ROI corresponding to “High” encompassed the ^15^N-positive signal of each OS and each “Low” ROI consisted of the distal ^15^N-negative OS section. Expression of *Ldhb* determined by IHC in a 30-day old *rd*^*1*^ retina. Glutamine synthetase (GS), a Mueller glia-specific marker, is used to demonstrate that the staining is within the cones, which are wrapped by Mueller glia cells. Panels on the right are higher magnification of boxed areas. ELM, external limiting membrane.

The data on glucose-derived ^13^C incorporation presented in Figure 1A and B are consistent with our previous finding that glycolysis plays a cell-autonomous role in photoreceptor anabolism^19^. Cones in the RP retina lose their OS prior to death, and recent evidence suggests that wild-type cones rely on aerobic glycolysis for maintenance of their OS and visual transduction (Rajala et al., 2018). This led us to hypothesize that, if glycolysis is inadequate, then augmentation of intracellular glycolytic intermediate(s) availability might improve the biosynthetic capacity of these cells and improve their function and/or survival. This was the rationale to examine the effect of promoting gluconeogenesis in cones.

Additional data indicated towards the feasibility of gluconeogenesis in RP cones. Previously we reported multiple lines of evidence that the LDHB subunit of lactate dehydrogenase is excluded from photoreceptors in a wild type retina (Chinchore et al., 2017). This isoenzyme is expressed in tissues that use lactate as a substrate for oxidative metabolism or gluconeogenesis (Dawson et al., 1964). LDHB catalyzes the equilibrium between lactate and pyruvate. Complete oxidation of pyruvate leads to release of its carbons as carbon dioxide. However, when the carbons are diverted, as happens in gluconeogenesis in the liver, the extracellular lactate-derived carbons are converted into intracellular glucose or glycogen. We observed that LDHB is detectable in RP cones after rod death (Figure 1C). Additionally, the *Ldhb* gene is reported to be regulated by oxygen in the chick retina (Buono and Lang, 1999). Thus, we speculated that its expression in RP cones might indicate a shift away from glycolytic metabolism towards a more oxidative phenotype and this altered metabolic environment might be conducive for gluconeogenesis.

Four enzymes are necessary for gluconeogenesis in humans (Supplementary figure 3). These essentially run the glycolytic pathway in reverse by catalyzing the bypass of the irreversible steps. We considered Glucose-6-phosphatase dispensable from the point of having cell-autonomous availability of glycolytic intermediates. We designed AAV vectors for the expression of the remaining 3 gluconeogenic genes and used the Cytomegalovirus (CMV) immediate-early enhancer and promoter (Figure 2A). This promoter drives strong expression in the cones in the mouse retina (Xiong et al., 2015). The vectors were tested for enzyme expression in 293T cells by plasmid transfection, as well as in retinas infected *in vivo* after packaging in the type 8 capsid (Figure 2B and 2C). In addition, the induction of glucose synthesis was assessed by enzymatic estimation of intracellular glucose in 293T cells transfected with plasmid vectors (Figure 2D). Although this method is not a measure of *in vivo* gluconeogenesis in cones, it allows an assessment of the constructs’ ability to promote glucose synthesis *in vitro*. This approach provides an estimation of glucose, as opposed to other hexoses, due to the specificity of the glucose oxidase used in the assay (see ****Methods****). This *in vitro* assay was necessary as it is technically challenging to maintain a degenerating retina in a metabolically stable state for conventional metabolomic flux analysis.

**Figure 2:**
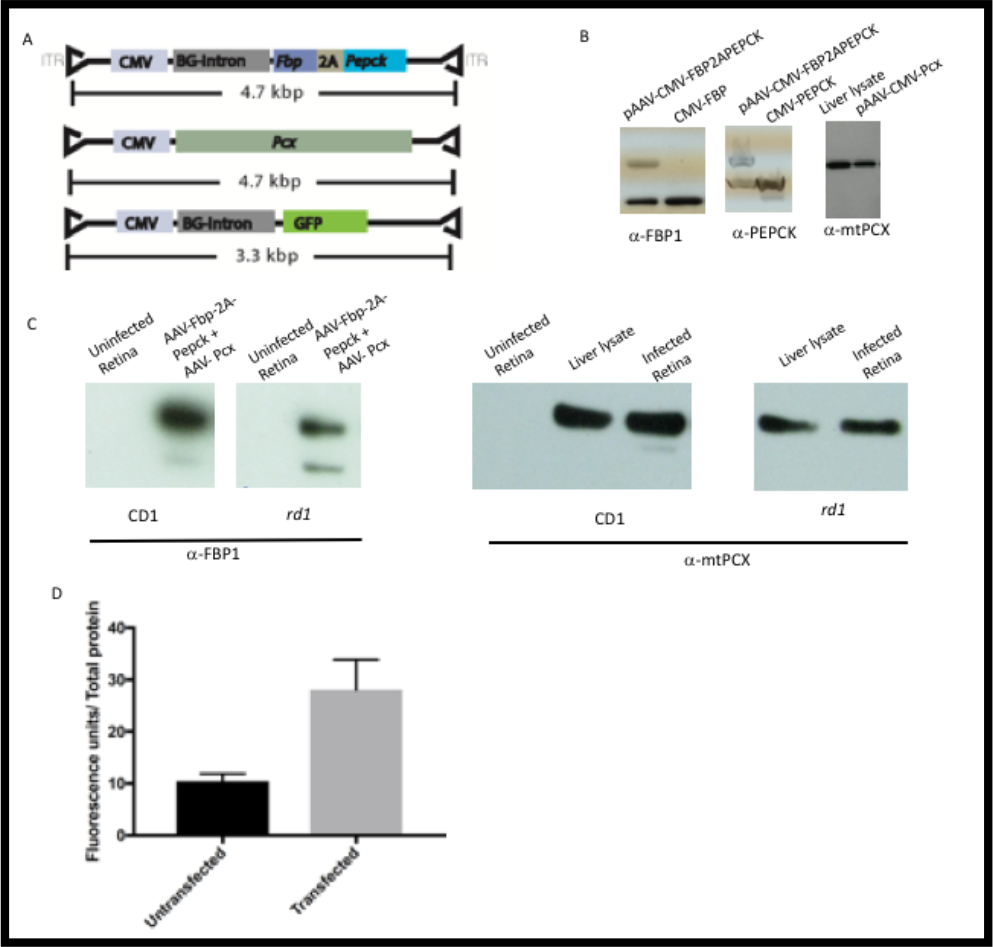
AAV vectors for expression of gluconeogenic enzymes. (A). Schematic representation of AAV-CMV-FBP-2A-PEPCK, AAV-CMV-PCX and, AAV-CMV-GFP constructs. BG-intron, human β-globin intron. ITR, inverted terminal repeats. (B). Western blot to assess protein expression from lysates prepared from HEK293T cells, transfected with constructs indicated on top of the lanes. Blots were probed with antisera indicated at the bottom of individual panels. The higher molecular weight band detected by anti-PEPCK and anti-FBP-1 is the calculated molecular weight of a fusion protein resulting from failure of the P2A sequence. (C). Western blot from lysates prepared from retinae of P25 wild type (CD1) or P29 *rd*^*1*^ mice transduced with AAVs indicated in panel A. Uninfected littermate retinae are used as controls. Blots were probed with antisera indicated at the bottom of the panel. (D). Glucose estimation from HEK293T cells cotransfected with plasmids used to generate AAV-CMV-Fbp-2A-Pepck and AAV-CMV-Pcx (Transfected). Transfected cells were incubated in glucose-deficient medium supplemented with sodium lactate for 48 hours. Untransfected cells subjected to identical medium changes as transfected cells were used as control (Untransfected). Data Mean±SD of three biological replicates.

Next, we assessed the effect of gluconeogenic gene expression on cone survival in *rd*^*1*^ mice. This strain of mice has a mutation in phosphodiesterase β (PDEβ), which leads to rapid rod death, followed by cone death (Figure 3A). The two AAV-gluconeogenesis vectors were coinjected with a third AAV encoding Green Fluorescent Protein (GFP), which enables tracking of infected cells. The three AAVs were mixed in a 1:1:1 ratio and injected into the subretinal space of neonatal *rd*^*1*^ mice. The photoreceptor degeneration in mice follows a center-to-periphery pattern, thus at a given time point, cells closer to the center are in a more advanced stage of disease than those in the periphery (Figure 3B). We quantified the cone density in the central and peripheral retina in animals infected with AAV vectors encoding gluconeogenic genes and GFP (gluconeogenic group). As a control, we injected only AAV-GFP, using an equivalent number of AAV-GFP genome copies (gc) as used for the gluconeogenic group (Figure 3C and 3D). Increased cone density was observed in the retinae of the gluconeogenic group compared with the GFP control at P58 in *rd*^*1*^ mice.

**Figure 3:**
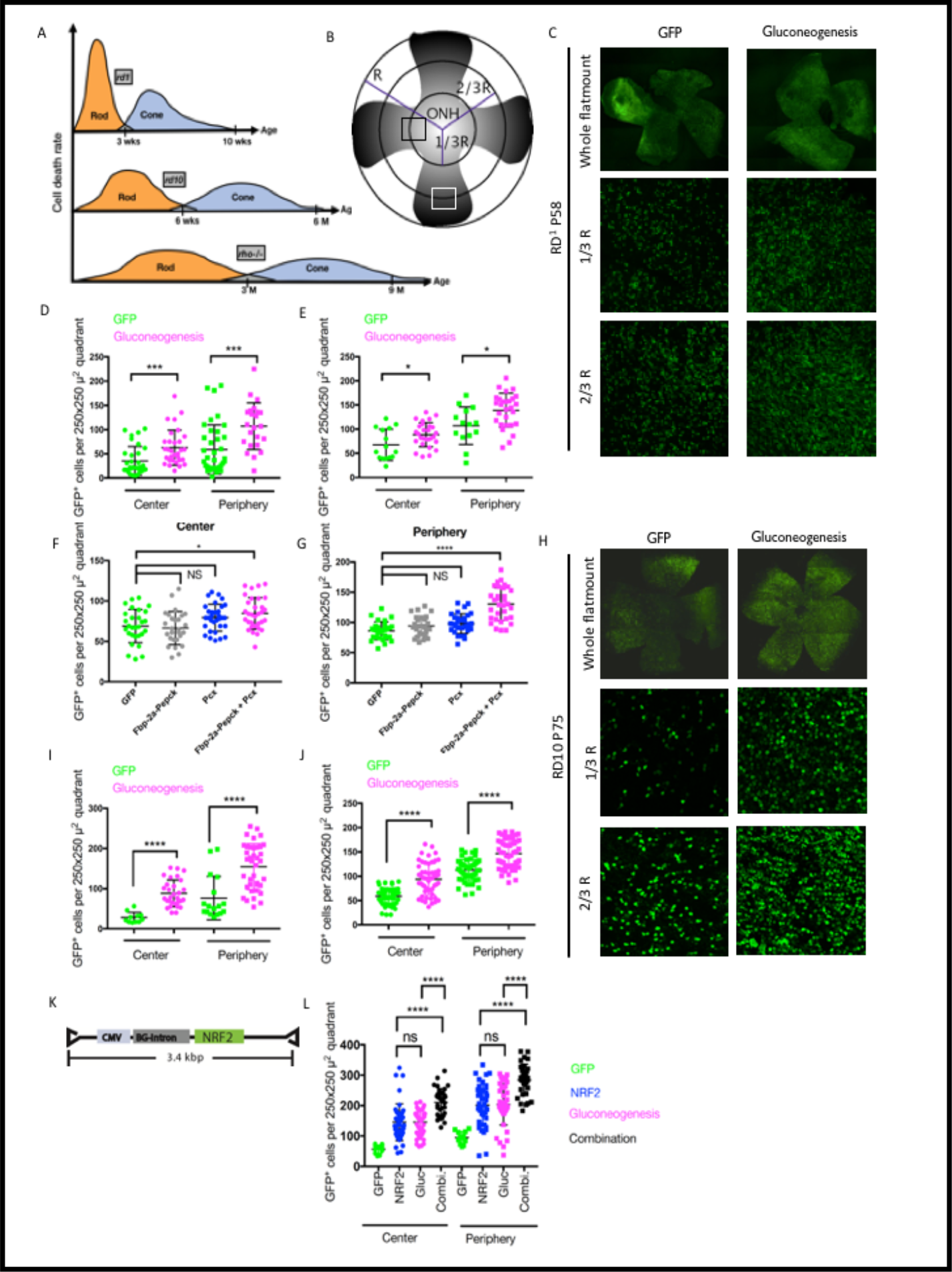
Effects of gluconeogenic gene expression on cone survival and additive effects with Nrf2. (A). Illustration of rod and cone death rates in the 3 RP models used in the current study. (B). Schematic of method of cone density quantification in a retinal flatmount. The gradient of surviving cones from center to periphery is depicted. The central GFP^+^ cones were quantified in a 250μm × 250 μm square (black) placed along 1/3 of the retinal radius (R) with the Optic Nerve Head (ONH) as the geographic center. Peripheral GFP^+^ cones were quantified in a 250μm × 250 μm square (white) placed along 2/3 of the retinal meridian. Cone densities in 3-4 such squares in the center and periphery were calculated for each well-infected retina. (C). Retinal flatmounts of P58 *rd*^*1*^ imaged for GFP. Animals were subretinally injected on P1 with type 8 encapsidated AAV-CMV-GFP (GFP) or AAV-CMV-GFP, AAV-CMV-Fbp-2A-Pepck and, AAV-CMV-Pcx (Gluconeogenesis). Higher magnifications of a 250um × 250 um region from center and periphery are depicted. (D). Cone density measured as surviving GFP^+^ cones in the center and periphery of P58 *rd*^*1*^ injected with type 8 AAVs, represented in panel C. Each data point represents GFP^+^ cells per 250um × 250 um square. AAV-CMV-GFP (GFP, green), (n=11 retinae). AAV-CMV-Fbp-2A-Pepck + AAV-CMV-Pcx+ AAV-CMV-GFP (Gluconeogenesis, magenta), (n=9 retinae). Mean±SD represented in error bars. P-value, Center= 0.0373, Periphery= 0.0162. Unpaired *t*-test with Welch’s correction. (E). Cone density measured as surviving GFP^+^ cones in the center and periphery of P45 *rd*^*1*^ injected with AAV encapsidated with Anc80L65. AAV-CMV-GFP (GFP, green), (n=5 retinae). AAV-CMV-Fbp-2A-Pepck+ AAV-CMV-Pcx+ AAV-CMV-GFP (Gluconeogenesis, magenta), (n=9 retinae). Mean±SD represented in error bars. Unpaired *t*-test with Welch’s correction. (F, G). Cone density measured as surviving GFP^+^ cones in the center (F) and periphery (G) of P45 *rd*^*1*^ injected with type 8 AAVs. P0-P2 animals were injected with AAV-CMV-GFP, either alone (GFP) (n=9) or with AAV-CMV-Fbp-2A-Pepck (n=8), AAV-CMV-Pcx (n=9) or with AAV-CMV-Fbp-2A-Pepck plus AAV-CMV-Pcx (n=9). Mean±SD represented in error bars. Significant difference among means calculated by one-way Anova with Tukey’s multiple comparison test. (H). Flat-mounted P75 *rd*^*10*^ retinae infected with AAV8-CMV-GFP (GFP) or AAV8-CMV-GFP, AAV8-CMV-Fbp-2A-Pepck and, AAV8-CMV-Pcx (Gluconeogenesis). Higher magnifications of a 250mm × 250 mm region from center and periphery are depicted. (I). Quantification of cone density in P75 *rd*^*10*^ retinae represented in panel H. AAV-CMV-GFP (GFP, green), 6 retinae. AAV-CMV-Fbp-2A-Pepck+ AAV-CMV-Pcx+ AAV-CMV-GFP (Gluconeogenesis, magenta), 11 retinae. Mean±SD represented in error bars. Unpaired *t*-test with Welch’s correction (Center) and Kolmogorov-Smirnov correction for normality (Periphery). (J). Quantification of cone density in P122-130 *Rho*^*−/−*^ retinae injected P0-P3 with type 8-encapsidated AAV-CMV-GFP (GFP, green) (n=14 retinae) or AAV-CMV-Fbp-2A-Pepck+ AAV-CMV-Pcx+ AAV-CMV-GFP (Gluconeogenesis, magenta), 11 retinae. Mean±SD represented in error bars. Unpaired *t*-test with Welch’s correction (Center) and Kolmogorov-Smirnov correction for normality (Periphery). (K). Schematic of the AAV construct used to express NRF2. (L). Cone density in P75 *rd*^*10*^ retinae injected with AAVs between P0-P3. Mice were injected either with AAV8-GFP alone (GFP, green) (n=4 retinae) or AAV8-GFP plus AAV8-NRF2 (NRF2, blue) (n=13 retinae), AAV8-GFP plus AAV8-CMV-Fbp-2A-Pepck and AAV8-CMV-Pcx (Gluconeogenesis, magenta) (n=12 retinae) or all the above (Combination, black) (n=10 retinae). Quantification from center (circles) and periphery (squares) was independently assessed for Gaussian distribution of data points. Error bars, Mean±SD. Statistics, Ordinary one-way Anova with Dunn’s multiple comparison (Center) and one-way Anova with Tukey’s correction (Periphery).

We also tested the ability of gluconeogenic genes to prolong cone survival when viral genomes were encapsidated with Anc80L65- a synthetic capsid that has similar tropism as serotype 8 (Zinn et al., 2015). Compared to the AAV-GFP control group, the gluconeogenic group had significantly higher cone survival in the center as-well-as peripheral retina (Figure 3E).

We were interested in whether all three gluconeogenic genes were necessary for the increase in cone survival. The results of this experiment could provide data regarding the mechanism of survival. In addition, if the results indicated that injection of fewer virions was sufficient for survival, a reduction in viral load might reduce induction of an immune response. However, reducing the number of gluconeogenic genes that were delivered led to a loss of the rescue (Figure 3F and 3G) in P45 *rd*^*1*^ mice. We also compared cone survival achieved by gluconeogenic gene expression by type 8 and Anc80L65 encapsidated vectors by examining densities in equivalent regions of *rd*^*1*^ mice at equivalent timepoint (P45) (Figure 3E, F and, G). We found that expression of these genes via transduction of two capsid types resulted in equivalent (no significant difference) cone survival.

As the gluconeogenic therapy was designed to be a general strategy for the many types of RP disease families, we tested it in two other RP models harboring different rod-specific mutations (Figure 3A). The *rd*^*10*^ mouse has a different mutation than the *rd*^*1*^ in the *Pde*β gene and has slower kinetics of rod and cone cell death (Chang et al., 2007). The homozygous rhodopsin knockout (*Rho*^*−/−*^) mice harbor a null allele of the *Rhodopsin* gene, the most commonly mutated gene in autosomal dominant RP, and have significantly slower kinetics of photoreceptor death relative to *rd*^*10*^. Central and peripheral cone density was quantified following subretinal injections into neonatal mice of these strains (Figure 3H, 3I and 3J). The gluconeogenesis group had significantly higher cone density than the controls.

It is likely that there are multiple stressors that affect cone survival in RP. Oxidized proteins, nucleic acids and lipids have been observed in RP retinas (Komeima et al., 2006). We previously targeted oxidative stress via AAV-mediated overexpression of NRF2- a transcription factor that upregulates the antioxidant response (Xiong et al., 2015). AAV8-NRF2 (Figure 3K) was effective in prolonging cone survival in the three mouse strains tested here. We thus combined AAV8-Nrf2 with the gluconeogenic viruses, and AAV8-GFP, injecting subretinally in neonatal *rd*^*10*^ mice (Figure 3L). The central and peripheral cone density was compared to those of cohorts that received either gluconeogenesis alone or NRF2 alone. These analyses indicated: (1). Gluconeogenesis was as effective as Nrf2 therapy in promoting cone survival and, (2). Combination of the two therapeutic approaches increased cone survival compared to either of the individual treatments.

During retinal degeneration, vision deteriorates before cone death, and the decline in vision is correlated with loss of outer and inner segments. Given the metabolic implication of gluconeogenic therapy and the relevance of the glycolytic pathway to outer segment maintenance, we evaluated the effect of this therapeutic intervention on the preservation of outer and inner segments. Retinae of *rd*^*1*^ mice transduced with AAVs encoding gluconeogenesis genes (and GFP), or GFP alone, were stained with Peanut Agglutinin (PNA), which labels the extracellular matrix surrounding the cone outer and inner segments (Figure 4A and 4B). Cones transduced with the gluconeogenic AAVs had significantly more labelling with PNA. An overall improvement in cone morphology also was noted in this group.

**Figure 4:**
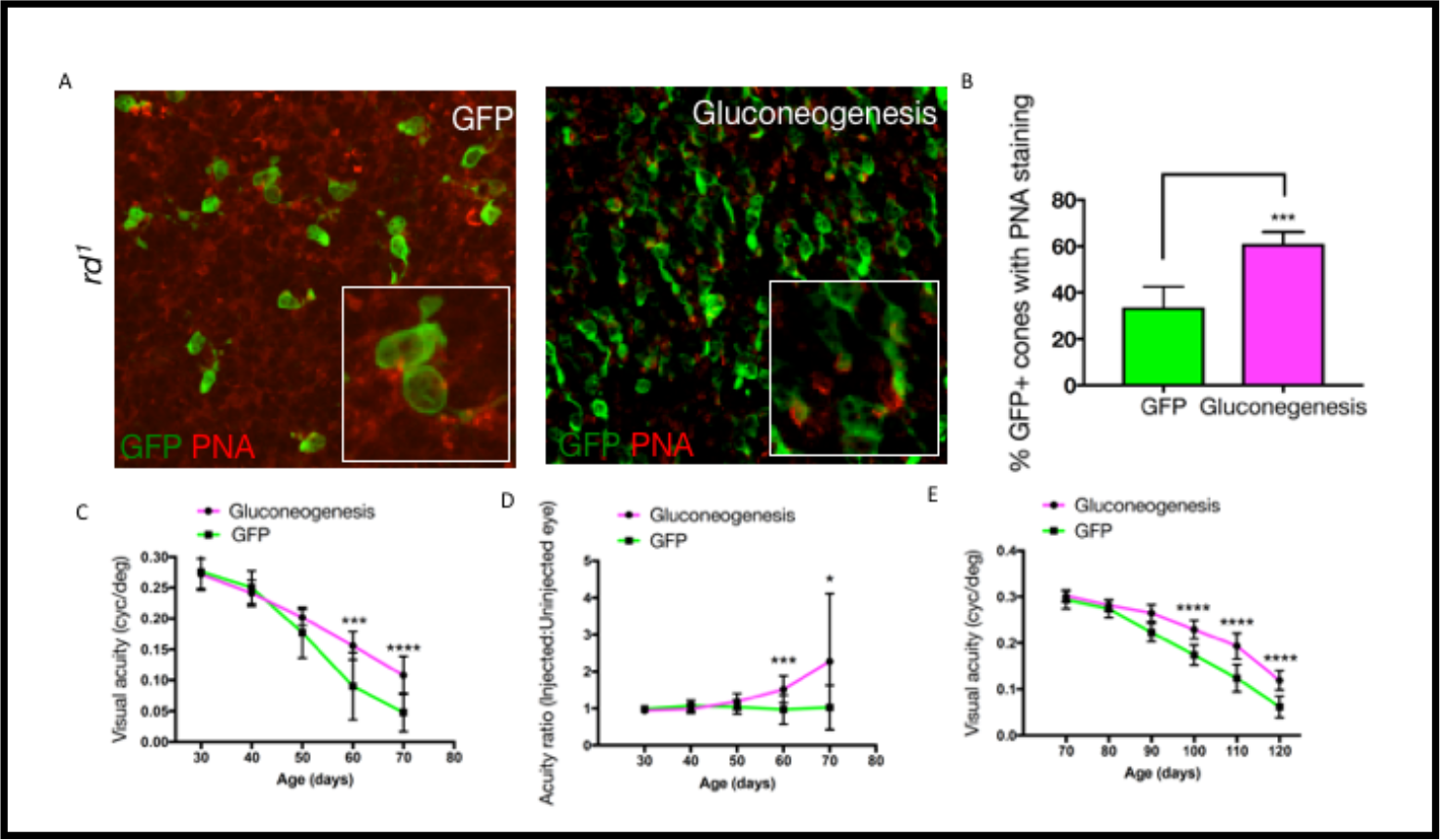
Effects of gluconeogenic gene expression on cone morphology and visual acuity in RP mice. (A). PNA staining of surviving cones in P45 *rd*^*1*^ retina infected at P1 with either AAV-CMV-GFP alone (GFP panel), or with AAV-CMV-Fbp-2A-Pepck and AAV-CMV-Pcx (Gluconeogenesis). Each panel is a 150 μm × 150 μm crop of original tile scan of an entire retina from mid-periphery. Inset, 3X higher magnification. (B). Quantification of GFP^+^ cells positive for PNA in P45 *rd*^*1*^ retina. Approximately 150 cells were counted per retina in each group from mid-periphery. Data, Mean±SD. Unpaired *t-*test with Welch’s correction. GFP, (n=6 retinae). Gluconeogenesis, (n=8 retinae). Optomotor assay to assess visual acuity of P0-P2 injected *rd*^*10*^ mice. Mice were injected in their right eyes with AAV-CMV-GFP alone (GFP) or AAV-CMV-GFP with AAV-CMV-Fbp-2A-Pepck and AAV-CMV-Pcx (Gluconeogenesis). The left eye was not injected but assayed for acuity. Error bars, Mean±SD. GFP, *n*= 15 mice P30-P60 and *n*=14 mice at P70. Gluconeogenesis, *n*=14 mice P30-P40 and *n*= 13 mice at P70. The ratio of same-animal right (injected) eye and left (uninjected) eye visual acuity in Gluconeogenesis and GFP groups as a function of age of *rd*^*10*^. Error bars, Mean±SD. Optomotor assay to assess visual acuity of P0-P3 injected *Rho*^*−/−*^ mice. Mice were injected with type 8 AAV-CMV-GFP alone (GFP) or AAV-CMV-GFP with AAV-CMV-Fbp-2A-Pepck and AAV-CMV-Pcx (Gluconeogenesis). Error bars Mean±SD. GFP, *n*= 12 mice. Gluconeogenesis, *n*=12 mice. Statistics, unpaired, two-tailed *t-*test with Welch’s correction for panels C, D and, E.

The goal of this therapy was not only to increase cone survival and morphology, but to prolong vision. To test if increased cone survival and improved cone morphology by gluconeogenesis correlated with better outcomes in vision, a longitudinal study to assess the visual acuity of infected mice was carried out. We evaluated the visual acuity by the optomotor assay, that measures the involuntary head movement in the direction of a mobile visual stimulus (Prusky et al., 2004). The right eyes of *rd*^*10*^ mice were injected with the AAVs encoding the gluconeogenic genes and GFP, or with GFP alone. The left eyes were uninjected. The visual acuity of left and right eye for each mouse was quantified and plotted either as a ratio of right to left, or without normalization, as a function of age (Figure 4C and D). The injected eyes that received gluconeogenesis genes performed significantly better than those that received AAV-GFP (Figure 4C). The visual acuity of the injected right eye of animals that received AAV-GFP was equivalent to that of the uninjected left eye (Figure 4D). In contrast, despite an age-dependent decline (Figure 4D), the visual acuity of the right eye of the gluconeogenesis group consistently outperformed the uninjected left eye, evident as an improved ratio (Figure 4D). We also compared the effect of gluconeogenesis on visual acuity in *Rho*^*−/−*^ eyes where the retina degenerates slowly over an extended period of time. Consistent with improved cone survival in this background, the gluconeogenesis group also exhibited improved visual acuity at each time point relative to AAV-GFP injected eyes. Thus, gluconeogenesis generically improves cone survival, morphology and function independent of the primary mutation afflicting the rods.

## Discussion

Here we show the utility of a gene therapy approach that uses gluconeogenic gene expression in cones to promote their survival in three different RP mouse models. Gluconeogenesis is an endergonic process. It is thus surprising that metabolically stressed cones not only tolerate it, but favorably respond to it. Among several possible scenarios include metabolic realignment to dampen a pathological metabolic pathway, increased usage of alternative metabolic fuels such as lactate or gluconeogenic amino acids, reinstatement of a favored intracellular metabolic environment or activation of signaling mechanisms that promote cell survival. The mechanistic basis for this rescue cannot be ascertained by the experiments performed in this study.

Together, the results presented here support an idea that a metabolic intervention utilizing gluconeogenic gene expression using AAV can be extended to many cases of RP, independent of rod-specific etiology, in order to ameliorate cone pathology and preserve their function. This and similar approaches should expedite therapeutic outcomes that broadly benefit patient populations of diverse genetic backgrounds and can serve as stop-gap measures before more precise and targeted therapies are approved for clinical use.

## Methods

### Animal husbandry

All animal handling and procedures adhered to the Harvard Medical School’s IACUC guidelines. The FVB inbred strain of mice harboring the *Pde6β*^*rd1*^ allele were purchased as timed pregnant mothers from Charles River Laboratories, Boston, MA. The *rd10* strain was obtained from Jackson Laboratories, Bar Harbor, ME. *Rho*^*−/−*^ strain, described previously (Lem et al., 1999), was provided by Janice Lem. The *rd10* and *Rho*^*−/−*^ mice were maintained in the HMS mouse colony. WT albino (CD-1) mice were obtained from Charles River Laboratories. The animals were housed in a room with temperature set at 25 °C and set to a 12-hour light and 12-hour dark cycle. The light intensity inside the cages varied 300 lx at a cage position closest to the light source and 0-3 lx at a position farthest from the light source. The injected mice were held on rack spaces where intracage light intensity ranged between ~145 to 250 lx.

### AAV Vector preparation

Mouse full-length cDNA clones for Fbp1, Pepck and Pcx were obtained from MGC/Dharmacon. Fbp1 and Pepck were amplified with primer sequences listed in Table 1, which also introduced 2A peptide sequence. The two amplicons were cloned into the AAV-MCS8 vector (Harvard Medical School DF/HCC DNA Resource Center) using Gibson isothermal assembly (Gibson et al., 2009). For subcloning of Pcx, the β-globin intron was removed from the vector to adjust for the vector size limitations.

**Table 1.**
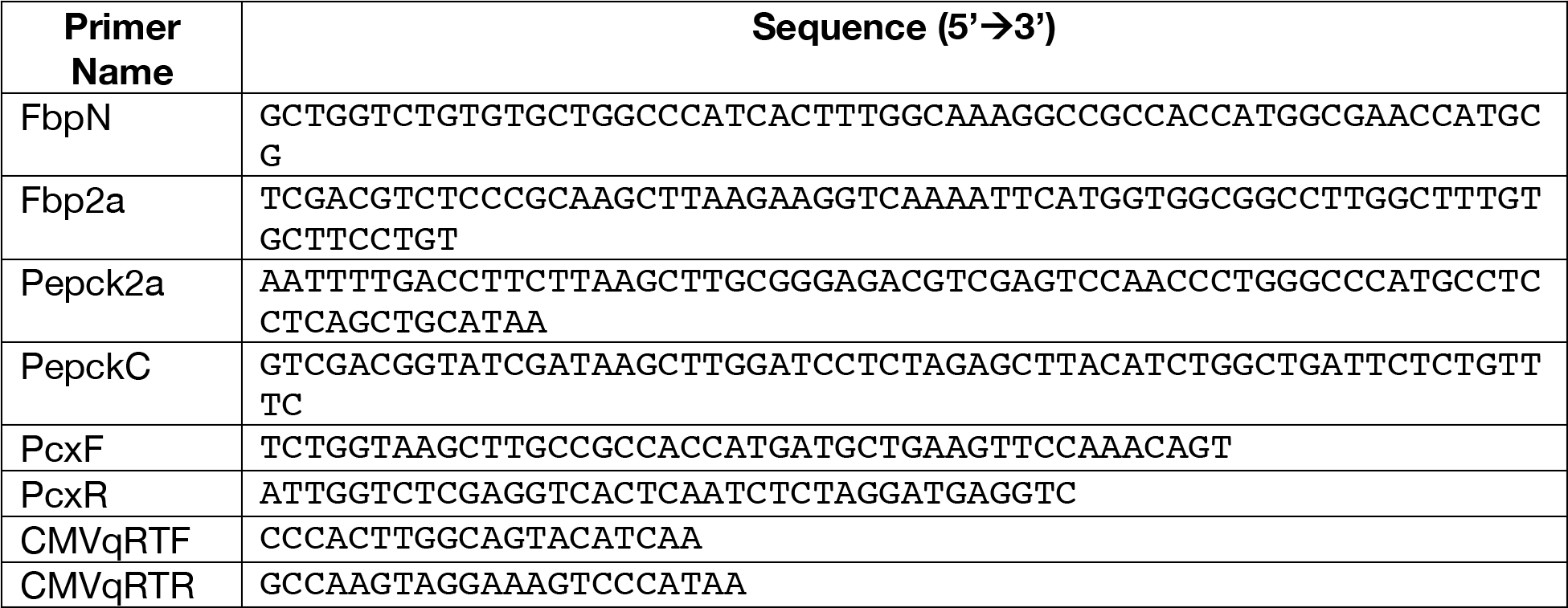

AAV8 viruses were prepared by Iodixanol density gradient centrifugation from the supernatant of HEK293T cells transfected with the viral vector and helper plasmids(Xiong et al., 2015). Additional batches of AAV8 and Anc80LC65 were prepared by Gene Transfer Core (Schepens Eye Institute, Boston, MA). The viruses were quantified by quantitative PCR using SYBR intercalating dye (Kapa) using primers specific for the CMV promoter region (Table 1) and a standard curve.

### Subretinal injections

The viruses were mixed to give a 1:1:1 ratio and mixed with fast green dye to aid visualization (Matsuda and Cepko, 2004). The identity of viruses was masked with decoded identifiers by laboratory colleagues who are not co-authors of this work. Approximately, 0.3 microliters of 1E12-1E13 gc/mL were injected subretinally in neonatal mouse pups using pulled glass needles controlled using Femtojet injector (Eppendorf).

### Glucose estimation

Twenty-four hours before transfection, ~4E6 HEK293T cells were seeded on a 10 cm plate. Three plates for gluconeogenic transfection groups and three plates for controls were seeded at the same time. The cells grew in Dulbecco’s modified Eagle’s medium (DMEM)- High Glucose (25mM) (Invitrogen, Catalog# 11965-118) with 10% Fetal Bovine Serum (FBS). Cells were transfected with plasmids with polyethylenimine (PEI) for ~16 hours in serum-free DMEM or were untransfected. The transfection medium was replaced with DMEM+5% FBS for 24 hours. The medium was then replaced with glucose and pyruvate-free DMEM (Invitrogen, Catalog# 11966-025) supplemented with Sodium Lactate (Sigma-Aldrich) to a final concentration of 15 mM and incubated for 36 hours. The medium was removed, and the cells were quickly rinsed with 1X Phosphate buffered saline (PBS) before addition of ice-cold hypotonic lysis buffer (Chinchore et al., 2017) without dithiothreitol, protease and phosphatase inhibitors to avoid interference in subsequent assays. The cells were subjected to one round of freezing and thawing on dry-ice:ethanol mixture and 37 °C followed by sonication. A fraction of lysate was preserved for protein estimation by Qubit assay (Invitrogen). The remainder lysate was centrifuged at 16,000×*g* at 4 °C for 30 minutes and supernatant rapidly transferred for glucose assay. The glucose estimation was carried out using Amplex Red Glucose assay kit (Invitrogen, Catalog# A22189) using previously unfrozen, freshly-resuspended glucose oxidase using excitation at 530 nm and fluorescence detection at 590 nm. Three technical replicates arising from each lysate were averaged. The three lysates from each condition (transfected and untransfected) served as biological replicates.

### Multi-isotope imaging mass spectrometry (MIMS)

Stable isotope tracers were purchased from Cambridge Isotope Laboratories. Tissue was fixed with 4% paraformaldehyde, embedded in EPON, sectioned to 0.5 microns, and mounted on silicon wafers. Samples were gold-coated and analyzed with the NanoSIMS 50L (CAMECA) at the Brigham and Women’s Hospital Center for NanoImaging as previously described (Guillermier et al., 2017a, 2017b; Kim et al., 2014; Steinhauser et al., 2012). ^2^H-labeling was quantified by measuring the ^12^C_2_^2^H^−^/^12^C_2_^1^H^−^ ratio. ^13^C-labeling was quantified by measuring the ^13^C^14^N^−^/^12^C^14^N^−^ ratio. ^15^N-labeling was quantified by measuring the ^12^C^15^N^−^/^12^C^14^N^−^ ratio. Images were viewed and processed, using a custom plugin to ImageJ (OpenMIMS 3.0: https://github.com/BWHCNI/OpenMIMS).

### Antibodies, reagents, IHC and Imaging

Antibodies against FBP1, PEPCK and PCX are listed in Table 2. Rhodamine-conjugated Peanut Agglutinin (PNA, Vector Laboratories) was used at 1:700. Retinae were dissected and fixed in 4% paraformaldehyde for 30 minutes, washed with 1XPBS+ 0.1% Tween20 (PBST) and incubated in anti-GFP antibody (Table 2) overnight at 4 °C. Next day, the retinae were washed with PBST followed by Alexa-488 (Invitrogen) or DyLight-conjugated (Jackson Immunoresearch) anti-rabbit or anti-chicken antibody (1:1000) for 1 hour. The retinae were mounted between glass coverslips and imaged from the photoreceptor side on Zeiss LSM 780 or 710 confocal microscope. Tile scans were obtained on a 10X air objective. Experimental groups were processed side-by-side and imaged on the same session with identical settings.

**Table 2:**
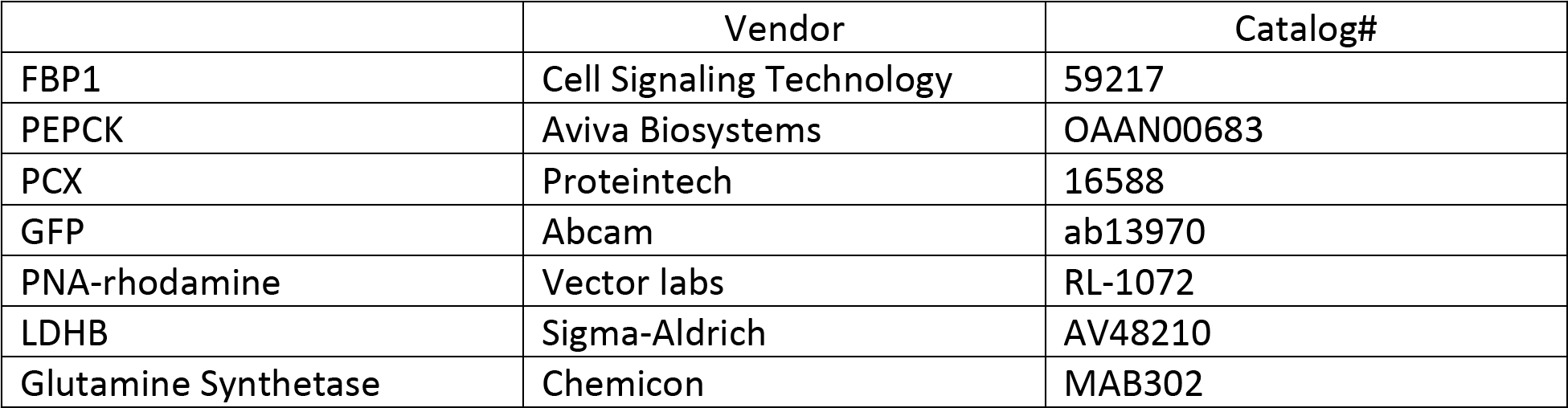

### Cone density quantitation

Images were imported on NIH ImageJ or FIJI. Each retinal flatmount image was processed individually. A circle was drawn with optic nerve head as center with the circumference touching the outer edges of retina. In a non-uniform retinal flatmount, the circumference of the circle touched at least two-thirds of the retinal edge. The radius (R) of the circle was then measured. Two concentric circles of 1/3R and 2/3R were then drawn. At least 3 squares of 250μm × 250 μm were drawn on the central and peripheral circles. The square was positioned such that the circumferential arc would bisect it as much as possible. The green GFP^+^ cones were manually counted per square. The number of cones in each square was used as a data point for each treatment group’s center or peripheral regions. All the data points were represented in the graphs to depict the entire range of transduced cones. The identity of treatment groups remained masked until quantification was complete. For samples where cone morphology was suggestive of a rescue, quantification was performed by at least two individuals. For, PNA labeling and quantification cones from region between the center (1/3R) and periphery (2/3R) were considered. This region was defined as mid-periphery.

### Visual acuity assessment

Visual acuity was measured using an Optometry System (CerebralMechanics) as described previously (Xiong et al., 2015). Animals were ear tagged for identification. All behavior analysis was performed between 9 AM-12 PM. Behavior analysis was performed by individuals who did not perform subretinal injections. The identity of treatment groups was masked from the individuals who performed subretinal injections as-well-as behavior analyses. After collection of optomotory data, the animals were sacrificed, and retinae were processed for imaging to assess the extent of infection. Data from only those animals which had full retinal infection were included in analysis. No other statistical test to identify outliers was applied. The treatment groups were unmasked only after analyses were completed.

### Statistics

All the treatment groups were represented in littermates as much as possible. The assignment of groups was done at random. The primary investigators were blinded to treatments in experiments involving assessment of cell survival and visual performance. No quantitative methods to predetermine sample size were employed. D’Augustino-Pearson omnibus test was used to determine normality. Non-parametric methods were used when data had non-Gaussian distribution. No methods to detect or eliminate outliers were used and all datapoints are included and depicted. *p-*value denoted as: Not significant (NS), p>0.05; *p≤0.05; **p≤0.01; ***p≤0.001; ****p≤0.0001.

## Acknowledgments

We thank Jonathan C. Tang and Ariel Aspiras for their help with double blind studies. Some datapoints in optomotor analyses were acquired by Parimal Rana, who was masked for their identity. He also verified cone density on key experiments. We gratefully acknowledge his technical help. We are thankful for the imaging support from Microscopy Resources on the North Quad (MicRoN) core at Harvard Medical School.

This work was supported by funding from Astellas, Howard Hughes Medical Institute and the National Institutes of Health grant R01 EY023291.

---

**Supplemental figure 1.**
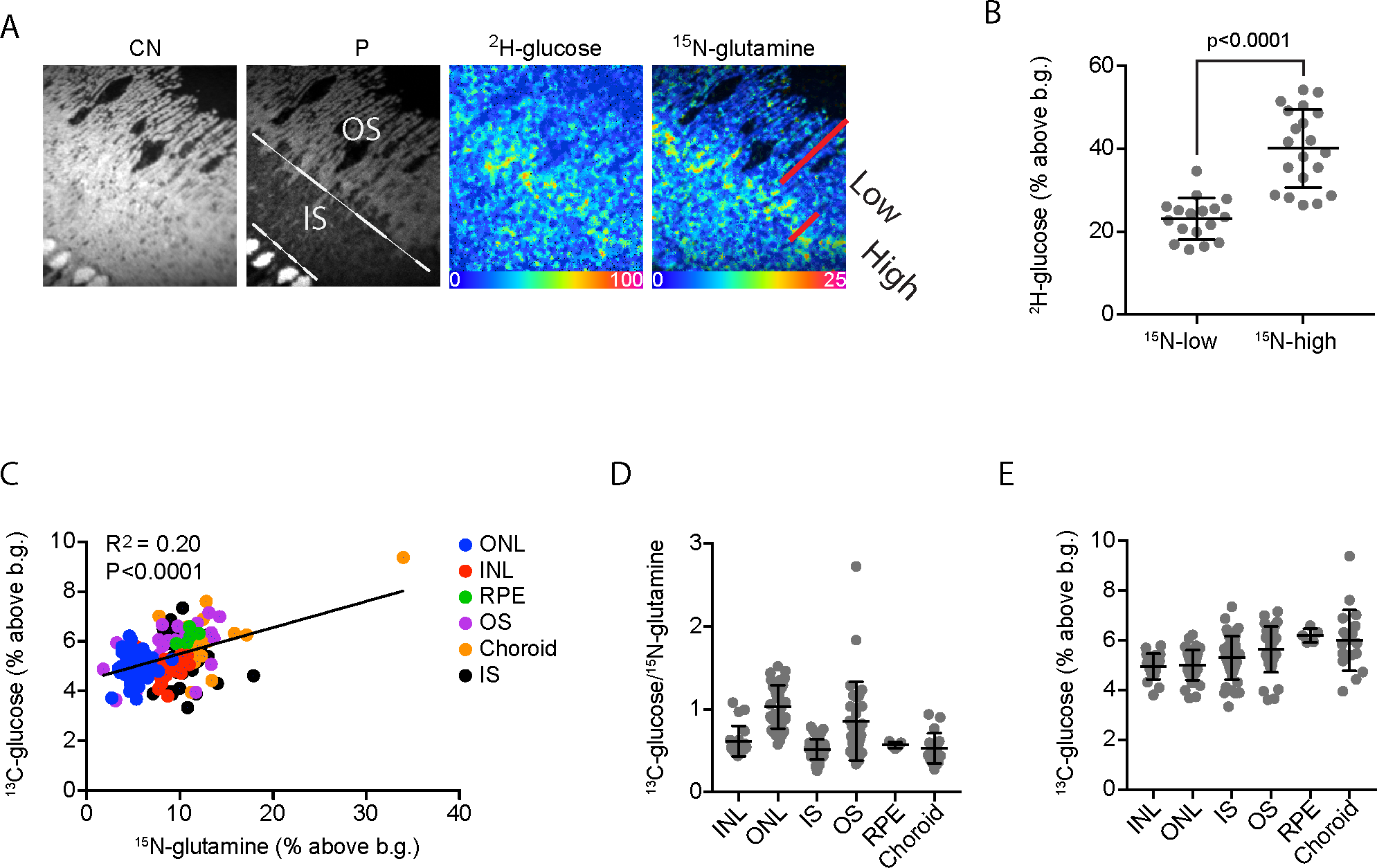
(A). MIMS imaging of an eye from wild type animal injected with ^2^H-glucose and ^15^N-glutamine. A scan from a region showing IS and OS is shown. Images derived from intensities of ^12^C^14^N (CN) and ^31^P (P) reveal stereotypical histological features of the eye. Incorporation of ^2^H-glucose and ^15^N-glutamine is shown visually by hue saturation intensity (HSI) ratio images with the blue end of the rainbow scale set to the natural background (0) and the upper bound of the scale is set at a % above natural background. (B) Quantification of ^2^H-glucose incorporation in regions of “High” or “Low” ^15^N-glutamine labelling. Individual region-of-interest (ROI) were selected with lateral dimensions not exceeding each OS width. A single ROI corresponding to “High” encompassed the ^15^N-positive signal of each OS and each “Low” ROI consisted of the distal ^15^N-negative OS section. (C). ^13^C-glucose incorporation as a function of ^15^N-glutamine. ROIs in indicated cells and subcellular structures were selected and quantified for ^13^C and ^15^N incorporation. Corresponding ^13^C labeling intensities were plotted as a function ^15^N incorporation. (D). Ratio of ^13^C:^15^N in selected regions of wild type eye. (E). Raw ^13^C-glucose incorporation values in selected regions in wild type eye.

**Supplemental figure 2.**
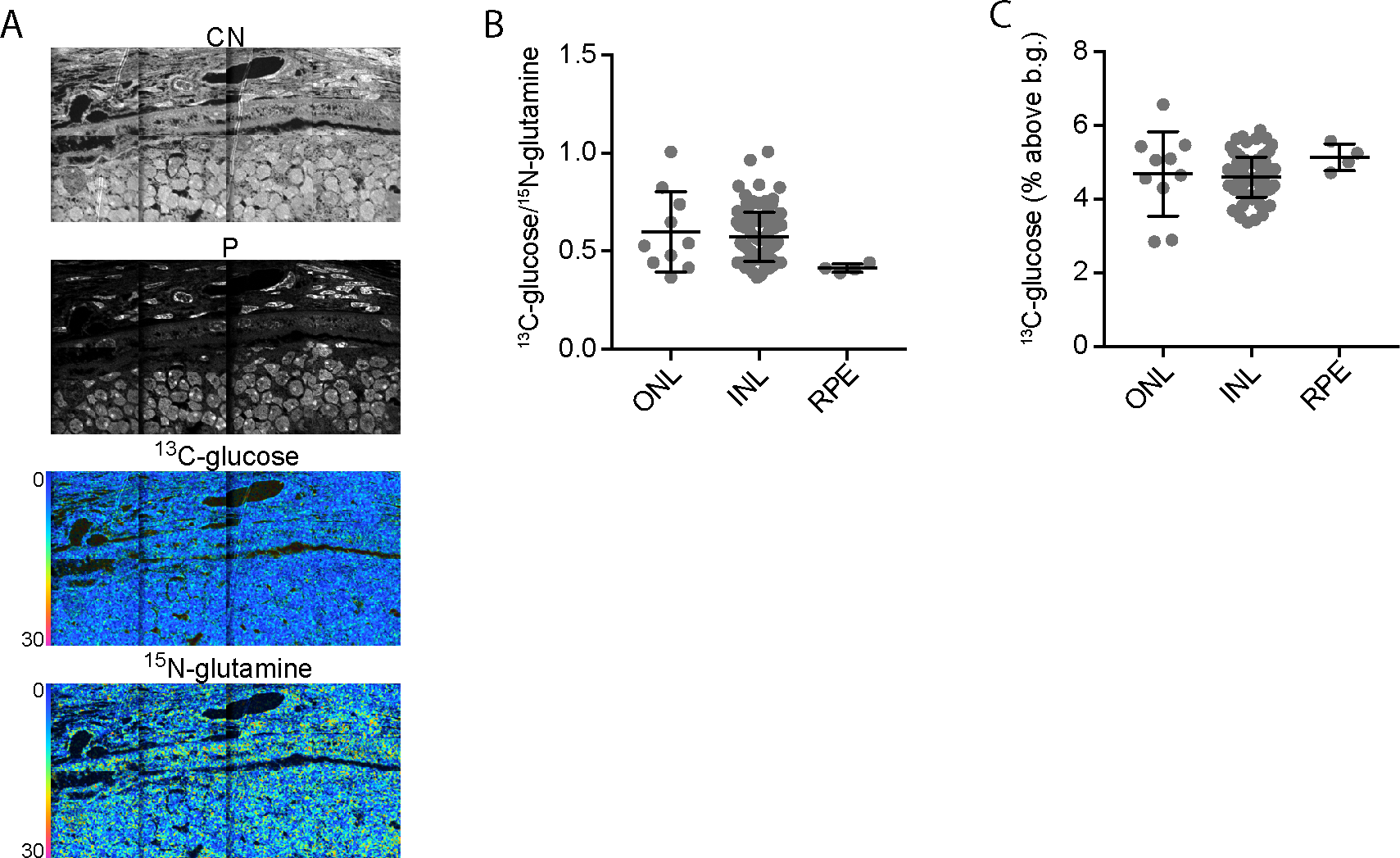
(A). MIMS imaging of an eye from 5-week old *rd*^*1*^ mouse injected with ^13^C-glucose and ^15^N-glutamine. (B). Ratio of ^13^C:^15^N in selected regions in *rd*^*1*^ eye. (C). Raw ^13^C-glucose incorporation values in selected regions in *rd*^*1*^.

**Supplementary figure 3:**
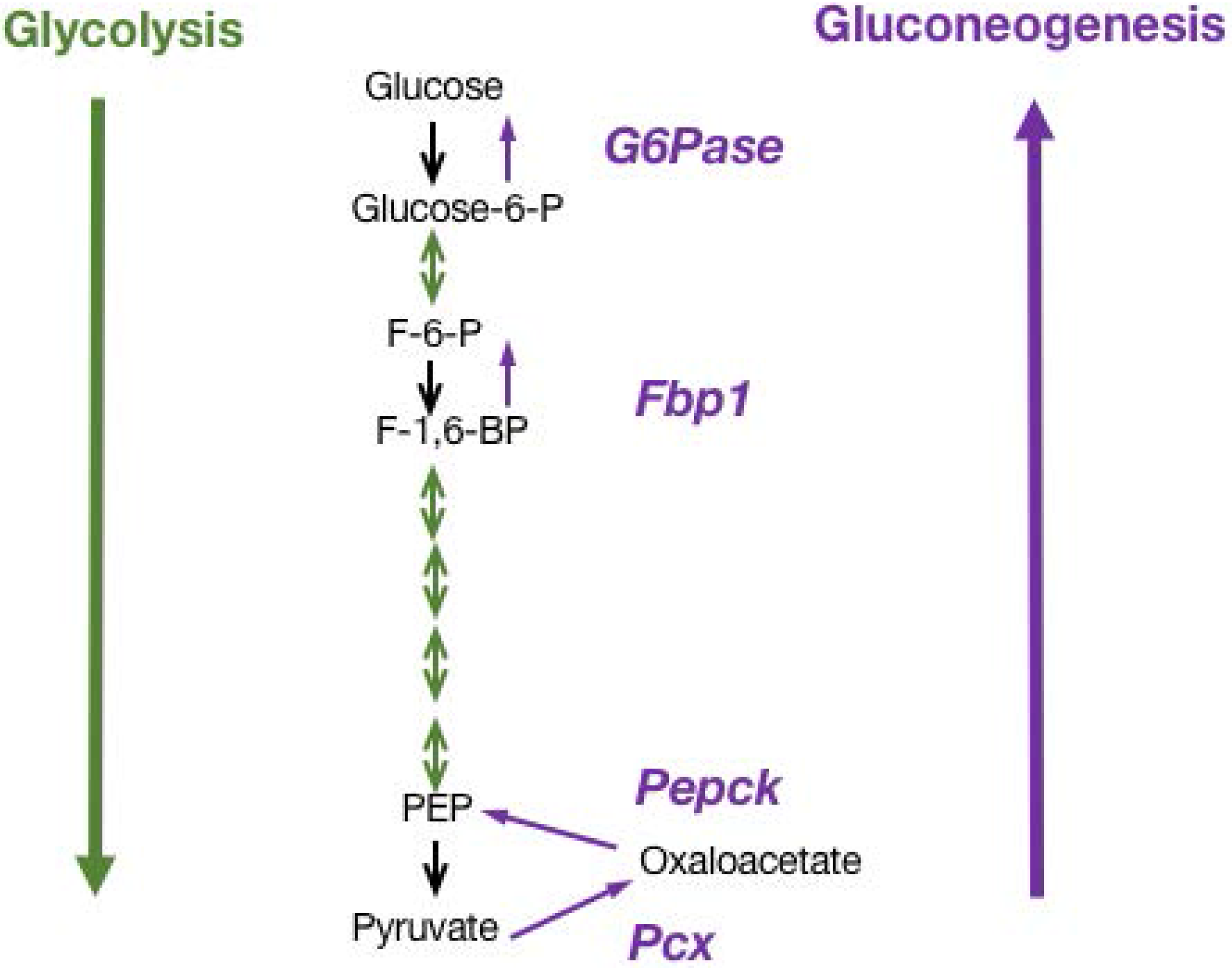
Schematic representation of glycolysis and gluconeogenic pathways. The key enzymes of gluconeogenesis are depicted. PEPCK, phosphoencolpyruvate carboxykinase. Pcx, pyruvate carboxylase. Fbp1, Fructose-1,6-bisphosphatase. G6Pase, glucose-6-phosphatase.

